# Revised subunit order of mammalian septin complexes explains their in vitro polymerization properties

**DOI:** 10.1101/569871

**Authors:** Forooz Soroor, Moshe S. Kim, Oliva Palander, Yadu Balachandran, Richard Collins, Samir Benlekbir, John Rubinstein, William S. Trimble

**Affiliations:** Cell Biology Program, Hospital for Sick Children, Toronto, Ontario, Canada, M5G 1X8; Department of Biochemistry, University of Toronto, Toronto, Ontario, Canada, M5G 1A8; Molecular Medicine Program, Hospital for Sick Children, Toronto, Ontario, Canada, M5G 1X8; Department of Physiology, University of Toronto, Toronto, Ontario, Canada, M5G 1A8

## Abstract

Septins are conserved GTP-binding cytoskeletal proteins that polymerize into filaments by end-to-end joining of heterooligomeric complexes. In human cells, both hexamers and octamers exist, and crystallography studies predicted the order of the hexamers to be SEPT7-SEPT6-SEPT2-SEPT2-SEPT6-SEPT7, while octamers are thought to have the same core, but with SEPT9 at the ends. However, based on this septin organization, octamers and hexamers would not be expected to co-polymerize due to incompatible ends. Here we isolated hexamers and octamers of specific composition from human cells and show that hexamers and octamers polymerize individually and, surprisingly, with each other. Binding of Borg3 results in distinctive clustering of each filament type. Moreover, we show that the organization of hexameric and octameric complexes is inverted compared to its original prediction. This revised septin organization is congruent with the organization and behavior of yeast septins suggesting that their properties are more conserved than was previously thought.

## INTRODUCTION

Septins are filamentous GTPases that were first discovered as genes important for cytokinesis in yeast (Hartwell et al., 1970). Septins are conserved from yeast to humans and the number of septins expressed is different between species. For example, *Caenorhabditis elegans* has just two (UNC-59 and UNC-61) while there are 13 in humans (SEPT1 through SEPT12 plus SEPT14) (Kinoshita, 2006; Beise et al., 2011). In mammals they play crucial roles in cytokinesis and many other processes such as cell polarity, spermatogenesis, exocytosis, phagocytosis, cell motility, ciliogenesis, and apoptosis (Fung et al, 2017). All septins possess a GTPase domain that belongs to the GTPase superclass of P-loop nucleotide triphosphatases, and the GTPase domain is flanked by variable N-and C-termini. Most septins have one or more coiled-coil extensions at the C-terminus involved in septin-septin interactions. Another conserved region near the N-terminus is a polybasic motif shown to bind to phosphoinositides (Zhang et al., 1999). Based on structural homology, it is possible to classify the 13 human septins into 4 subfamilies represented by their original founding members: SEPT2 subfamily (SEPTs 1,2,4,5); SEPT3 subfamily (SEPTs 3,9,12); SEPT6 subfamily (SEPTs 6,8,10,11,14); and SEPT7 subfamily (SEPT7 alone).

Septin heterooligomeric complexes, which consist of four, six or eight subunits, are building blocks for septin filament formation in different organisms. The well-studied yeast *Saccharomyces cerevisiae* septin complex (composed of Cdc3, Cdc10, Cdc11, and Cdc12) was isolated and shown to form filaments *in vitro* (Frazier et al., 1998). The yeast septin complex is a rod-shaped octamer and Bertin and colleagues (Bertin et al., 2008) revealed the order of yeast septins in the complex to be Cdc11-Cdc12-Cdc3-Cdc10-Cdc10-Cdc3-Cdc12-Cdc11 by antibody labeling or by attaching bulky tags such as maltose binding protein (MBP) or GFP to specific septins to visualize their position by electron microscopy (EM).

In mammals, septin complexes are likely to be more complicated given the larger number of genes expressed, but the presence of 4 subfamilies has led to speculation that mammalian septins may also exist in ordered complexes comprised of two copies of one member of each subfamily (Kinoshita, 2003). Blue native gels of HeLa cell lysates revealed that in these cells septins exist in two forms, hexamers (containing SEPT2, SEPT6, and SEPT7) and octamers (containing SEPT2, SEPT6, SEPT7 and SEPT9) (Sellin et al., 2014). Although not directly orthologous, the mammalian septins share structural traits with the yeast septins. For example, the SEPT3 subfamily lacks the coiled-coil in the C-terminal extension and the same is true for Cdc10 in yeast (Cao et al., 2007).

By expressing SEPT2, SEPT6 and SEPT7 in bacteria, Sirajuddin and colleagues (Sirajuddin et al., 2007) resolved the crystal structure of a human septin hexamer and determined the order of septins to be SEPT7-SEPT6-SEPT2-SEPT2-SEPT6-SEPT7. Unfortunately, the octameric complex containing SEPT9 is not easily expressed in bacteria. However, it was shown that SEPT9 associates with the complex by direct interaction with SEPT7 (Kim et al., 2011), implying a position at the ends of octamers. The crystal structure also revealed that septins in the unit complex alternately interacted via their NC (amino and carboxyl termini) and G (nucleotide-binding site) interfaces.

Intriguingly, the crystal structure prediction leads to two challenges for understanding septin biology in mammals. First, the most closely similar septins between yeast and mammals are not located in the same place in the complex. SEPT9 is structurally similar to Cdc10, while SEPT2 in more analogous to Cdc11, and this model places them at opposite positions in an octamer. Second, this organization would mean that octamers, which would end with the NC interface of SEPT9, would be unable to interact with hexamers, which end with the G interface of SEPT7. This would imply the presence of two sets of septin filaments in mammalian cells.

Given the predicted position of the SEPT3 subgroup at the terminal position in the octameric complex, the N and C termini would constitute the interaction surface involved in octameric septin polymerization. Since there are many SEPT9 isoforms that differ at their N and C termini, this raises the possibility that SEPT9 isoforms might influence octameric filament formation.

To test the role of different SEPT9 isoforms in septin octamer polymerization, and to compare the polymerization properties of octamers and hexamers, we established stable cell lines expressing defined septin complexes, purified the complexes, and used light and electron microscopy to monitor polymerization. Through these studies we provide surprising new insights into the organization of the septin complexes that challenge the existing models and demonstrate that the organization of septin complexes is structurally conserved between yeast and humans.

## RESULTS

### Purification and polymerization of defined mammalian septin complexes

Taking advantage of an inducible Flp-In^TM^ HeLa cell line expression system, we generated stable cell lines that can be induced to express either GFP-FLAG-His6-SEPT7 or GFP-FLAG-His6-SEPT9_v4 in HeLa cells. We confirmed the expression of the transgenes by western blotting using SEPT7, SEPT9, GFP or FLAG specific antibodies (Figure 1A) and show that they give rise to filamentous structures in the cell (Figure 1B).

**Figure 1:**
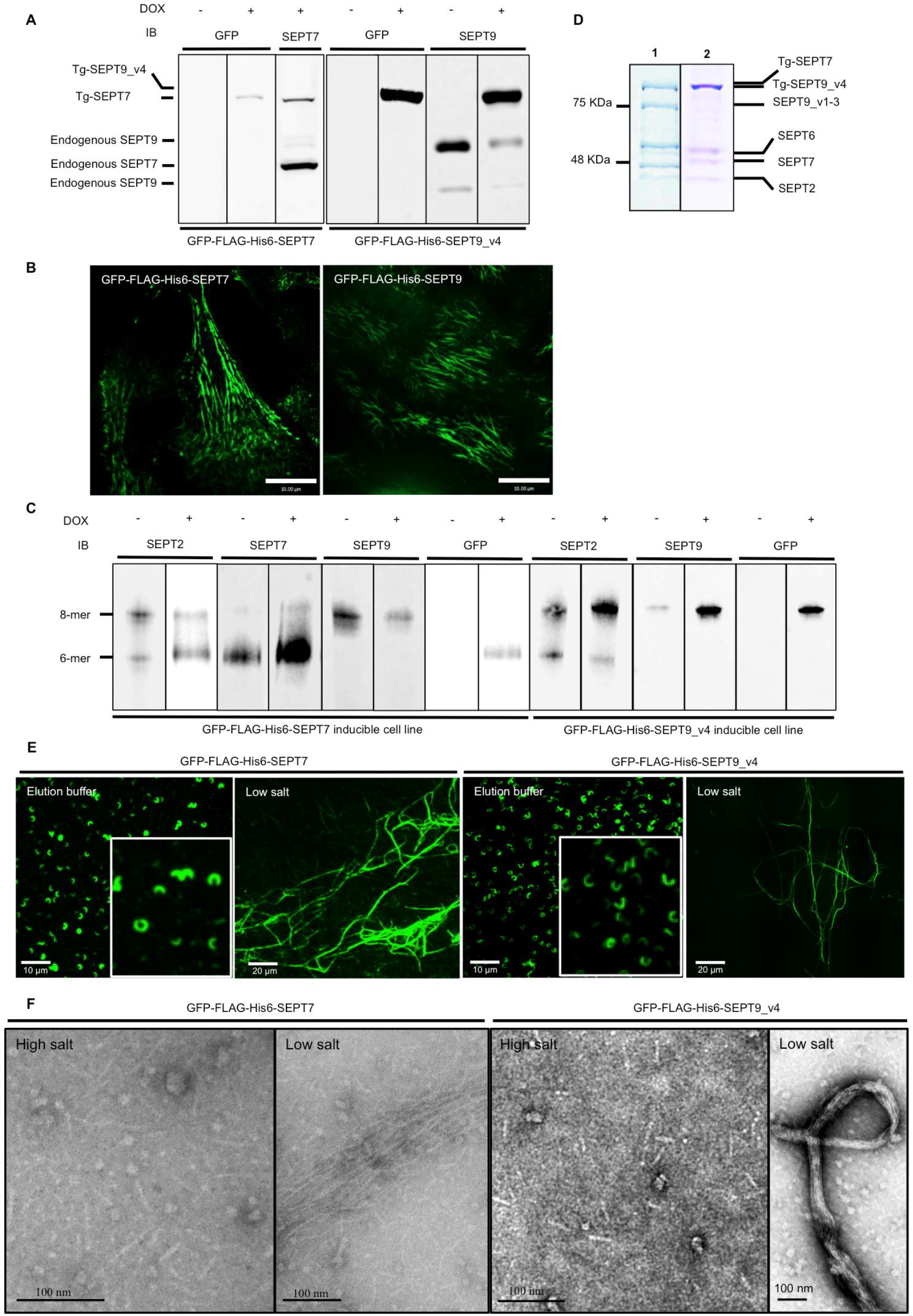
Expressed GFP-FLAG-His6-SEPT7 and GFP-FLAG-His6-SEPT9_v3 incorporate into endogenous septin filaments. (A) Western Blot of induced GFP-FLAG-His6-SEPT7 stable cell line lysate shows two bands (endogenous and transgene (Tg-SEPT7) when probed with anti-SEPT7 antibody and one band (Tg-SEPT7) when probed with anti-GFP antibody. Hexamer band containing transgene slightly larger than endogenous due to presence of tags. Induced GFP-FLAG-His6-SEPT9_v4 stable cell line lysate immunoblotted with SEPT9 and GFP antibodies shows three and one bands (endogenous and Tg-Sept9_v4) respectively in induced cells. Endogenous SEPT9 has two bands in uninduced cells representing different isoforms. (B) SEPT9 (GFP signal) is filamentous in cells, demonstrating expressed septins associated with stress fibers. (C) Septin heteromers of GFP-FLAG-His6-SEPT7 and GFP-FLAG-His6-SEPT9_v4 induced and uninduced stable cell lines resolved with blue native PAGE followed by immunodetection with SEPT2, SEPT7, and SEPT9 and GFP antibodies. Positions of hexamers and octamers determined previously (Sellin et al., 2014) (D) FLAG-immunopurified (1) GFP-FLAG-His6-SEPT7 and (2) GFP-FLAG-His6-SEPT9_v4 complex separated by SDS PAGE and stained with Coomassie blue. Protein position determined by Western blotting with specific antibodies (Fig S1) (E) Isolated complexes visualized by fluorescence microscopy microscopy in elution buffer (150 mM KCl) or following dialysis in low salt (50mM KCl). (F) Electron microscopy of isolated complexes in high salt (400 mM KCl) and low salt buffer (50mM KCl) following negative staining with uranyl acetate.

Sellin and colleagues (Sellin et al., 2011) used blue native gels to show that in mammalian cells, septin complexes are hetero-oligomeric hexamers or octamers. Blue native PAGE separates protein complexes based on their size and shape, and they found that septin hexamers migrate on the gel near the position of a 450 kDa globular protein marker while octamers run slower and have varied mobility depending on the size of SEPT3 family members or different SEPT9 isoforms expressed. Using the same protocol, we found that indeed there are both endogenous hexamers and octamers in the Flp-In Hela cell line (Figure 1C). Upon induction of the transgene GFP-FLAG-His6-SEPT7 or GFP-FLAG-His6-SEPT9_v4 we noted an increase in the relative proportion and intensity of the hexamer or octamer band respectively compared to non-induced cells (Figure 1C). This result confirmed that the expressed septin could interact with endogenous septins to form complexes. In addition, the expressed transgene typically caused the downregulation of the endogenous counterpart (Figure 1A), as has been reported previously (Sellin et al., 2014), likely due to competition for incorporation into the complexes.

Anti-FLAG immunopurification was performed following induction of GFP-FLAG-His6-SEPT7 or GFP-FLAG-His6-SEPT9_v4 for 24 hrs. Septin complexes were isolated and detected by Coomassie Blue staining after SDS-PAGE (Figure 1D) and individual bands could be detected at the size of each septin expressed in the cell. The identity of the endogenous septins in the purified complex was confirmed using antibodies specific for each septin, revealing the presence of SEPT2, SEPT6, SEPT7, and SEPT9 (Figure S1A) and indicating that the transgenes associated with endogenous septins in *bona fide* complexes. As expected, no complexes could be purified from cell lines that were not induced (data not shown).

Septin polymerization is affected by electrostatic interactions at the end subunits of the complex, and while polymerization is supported in low salt concentrations, it is inhibited in high salt (Frazier et al., 1998). Booth and colleagues (Booth et al., 2015) used a FRET-based system to show that full polymerization is supported at salt concentrations below 100 mM, and fully inhibited at concentrations above 300 mM with an EC50 of about 180 mM. Blue native gels were run at 450 mM KCl, ensuring full dissociation of the septin complexes and isolated complexes from GFP-FLAG-His6-SEPT9_v4 and GFP-FLAG-His6-SEPT7 cell lines run on the Blue Native PAGE were indeed octamers and hexamers (Figure S1B). Using negative staining electron microscopy, we measured the lengths of all complexes in high salt (400 mM). Septin rods containing SEPT9_v4 were 32-35 in length, consistent with the predicted length of octamers (Figure S1C and Sirajuddin et al., 2007). However, the size of complexes isolated from GFP-FLAG-His6-SEPT7 cell line fall mainly in the 25nm range (Figure S1C) expected for hexamers (Sirajuddin et al., 2007).

Mammalian septins localize along actin stress fibers in interphase cells and in the presence of actin-perturbing drugs the septins form rings with relatively uniform diameters (Kinoshita et al, 2002). Using the stable cell lines after cytochalasin D (CD) treatment and immunofluorescence, we observed septin rings and curved structures and found that GFP-FLAG-His-SEPT9 isoforms localized to these structures (Kim et al., 2011 and Figure S3). Since the protein is GFP-tagged, it enabled us to use fluorescence microscopy to visualize isolated septin complexes and filaments. Under elution conditions, the complexes were isolated in salt conditions (150mM KCl) predicted to support some polymerization. Therefore, isolated septins were spotted on coverslips in elution buffer and imaged by fluorescence microscopy. Interestingly, we observed curvilinear structures, spirals and rings *in vitro* that resemble those observed in the cells after CD treatment (Figure 1E and Figure S3). Upon dialysis of octamers or hexamers into low salt buffer (50mM KCl) we observed thin filaments, thick bundled filaments, rings and half rings using fluorescence microscopy (Figure 1E) and electron microscopy (Figure 1F).

### SEPT3 family members do not regulate polymerization

In yeast Cdc11 can be substituted at the ends of octamers with Shs1, leading to a significant change in septin polymerization from linear filaments to ring and gauze-like structures (Garcia et al., 2011) indicating the importance of the terminal septins on the polymerization process. In the case of mammalian septin octamers, if SEPT9 is located at the terminal position, then SEPT9-SEPT9 interactions would be critical for polymerization. Moreover, SEPT9 undergoes alternative splicing with five N-terminal splice variants (Peterson et al., 2010) raising the possibility that different isoforms of SEPT9 could have different functions. In support of this, we previously found that rescue of SEPT9 depleted cells with SEPT9_v3 could support cytokinesis, but SEPT9_v4 could not (Estey et al., 2010).

To determine whether octamers with different SEPT9 isoforms are all capable of forming filaments, the five major human SEPT9 variants were cloned into an inducible FRT vector inducible stable cell lines were generated individually expressing GFP-FLAG-His6-SEPT9_v1 through _v5. After characterizing these cell lines in denaturing and native conditions (Figure 2Ai and 2Bi), immunopurification using anti-FLAG antibodies resulted in the successful isolation of distinct septin complexes, as confirmed by Coomassie blue staining after SDS-PAGE and western blotting (Figure 2C and S2). Isolated complexes from these cell lines were spotted on coverslips and imaged by fluorescence microscopy. Septin rings, spirals, and curvilinear structures were observed in elution buffer by fluorescence microscopy (Figure 3A). Isolated septin complexes were supplemented with high salt buffer (400 mM KCl) to ensure that all septin higher order structures were disassembled to septin complexes. Negative staining electron microscopy confirmed that complexes isolated from all five stable cell lines are rod-shaped octamers (Figure 3C and S3). To further confirm that isolated complexes are octamers, purified septin complexes were run alongside cell lysates on blue native PAGE. The position of the bands on blue native PAGE in isolated complexes was identical to those seen in the lysates (Figure S2).

**Figure 2:**
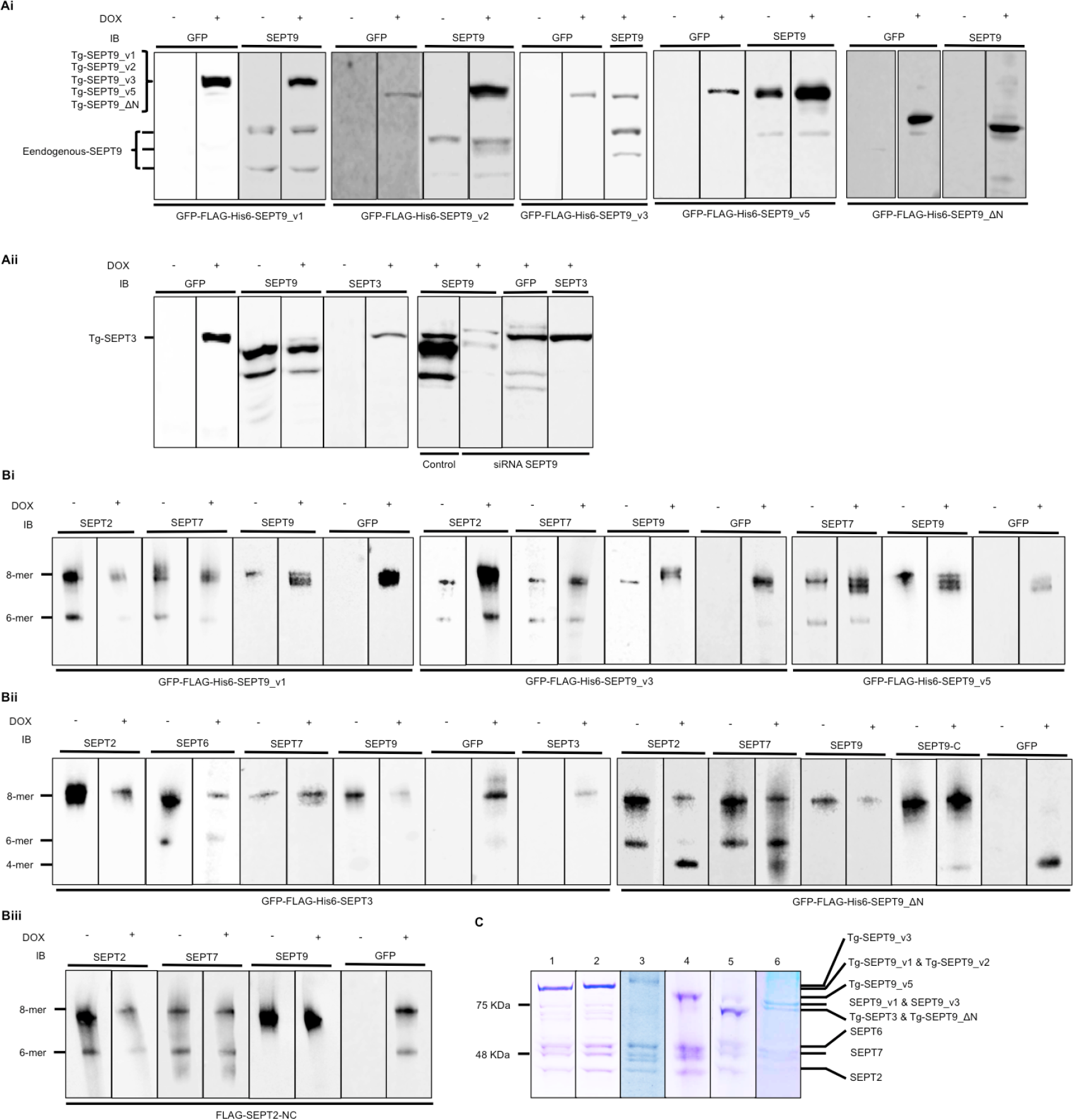
Generation of stable cell lines characterization. (Ai) Induced expression of GFP-FLAG-His6-SEPT9_v1 through _v5 in Flp-In HeLa cells immunoblotted with SEPT9 and GFP antibodies. In the case of GFP-FLAG-His6-SEPT9_v5, which is one of the shortest SEPT9 isoforms, co-migrates with the longest endogenous SEPT9. (Aii) Expression of GFP-FLAG-His6 SEPT3 was confirmed with immunoblotting with GFP, SEPT9 and SEPT3 antibodies. Endogenous SEPT9 was levels are diminished upon SEPT3 induction (Left panels). siRNA depletion of SEPT9 significantly reduces SEPT9 levels compared to SEPT3 (right panels). (Bi) Separation of septin complexes by blue native PAGE. GFP-FLAG-His6-SEPT9_v1 expression caused an increase in the thickness of the octameric band when compared to uninduced band immunoblotted for SEPT2, SEPT6, SEPT7, and SEPT9. Three separate bands in some SEPT9 blot may represent different octamers containing 0, 1 or 2 transgenes. GFP blot reveals presence of a transgene primarily band corresponding to the size of the transgene – for example GFP-FLAG-His6-SEPT9_v5 predominantly appears in the smallest of the three SEPT9 bands. Immunoblotting for SEPT2 antibody clearly shows an increase in the octameric band relative to hexamers following SEPT9 induction. (Bii) The lysate of expressed GFP-FLAG-His6-SEPT3 and GFP-FLAG-His6-SEPT9_ΔN along with uninduced cell lysates was resolved with blue native PAGE and immunoblotted for SEPT2, SEPT6, SEPT7, SEPT9, SEPT3, and GFP antibodies to show the participation of expressed septins into endogenous heterooctamers. Note that SEPT9_ΔN results in the formation of a higher mobility species below the hexamer band position that contains SEPT2, SEPT7 and GFP. (Biii) 3xFLAG-SEPT2-NC inducible stable cell line by blue native PAGE followed by immunodetection with SEPT2, SEPT7, SEPT9 and FLAG antibodies indicates 3xFLAG-SEPT2-NC-containing complexes are hexamers and octamers. (C) Lanes 1-6 represent FLAG-immunopurification of septin complexes from induced stable cell lines; (1) GFP-FLAG-His6-SEPT9_v1; (2) GFP-FLAG-His6-SEPT9_v2; (3) GFP-FLAG-His6-SEPT9_v3; (4) GFP-FLAG-His6-SEPT9_v5; (5) GFP-FLAG-His6-SEPT9_ΔN; and (6) GFP-FLAG-His6-SEPT3. Coomassie blue–stained SDS–PAGE gel indicates purity of septin complexes isolated from stable cell lines. Since a part of FLAG tag sequence (DYKDDDDK) interacts strongly with Coomassie blue, the expressed septin band appears darker than endogenous septin bands.

**Figure 3:**
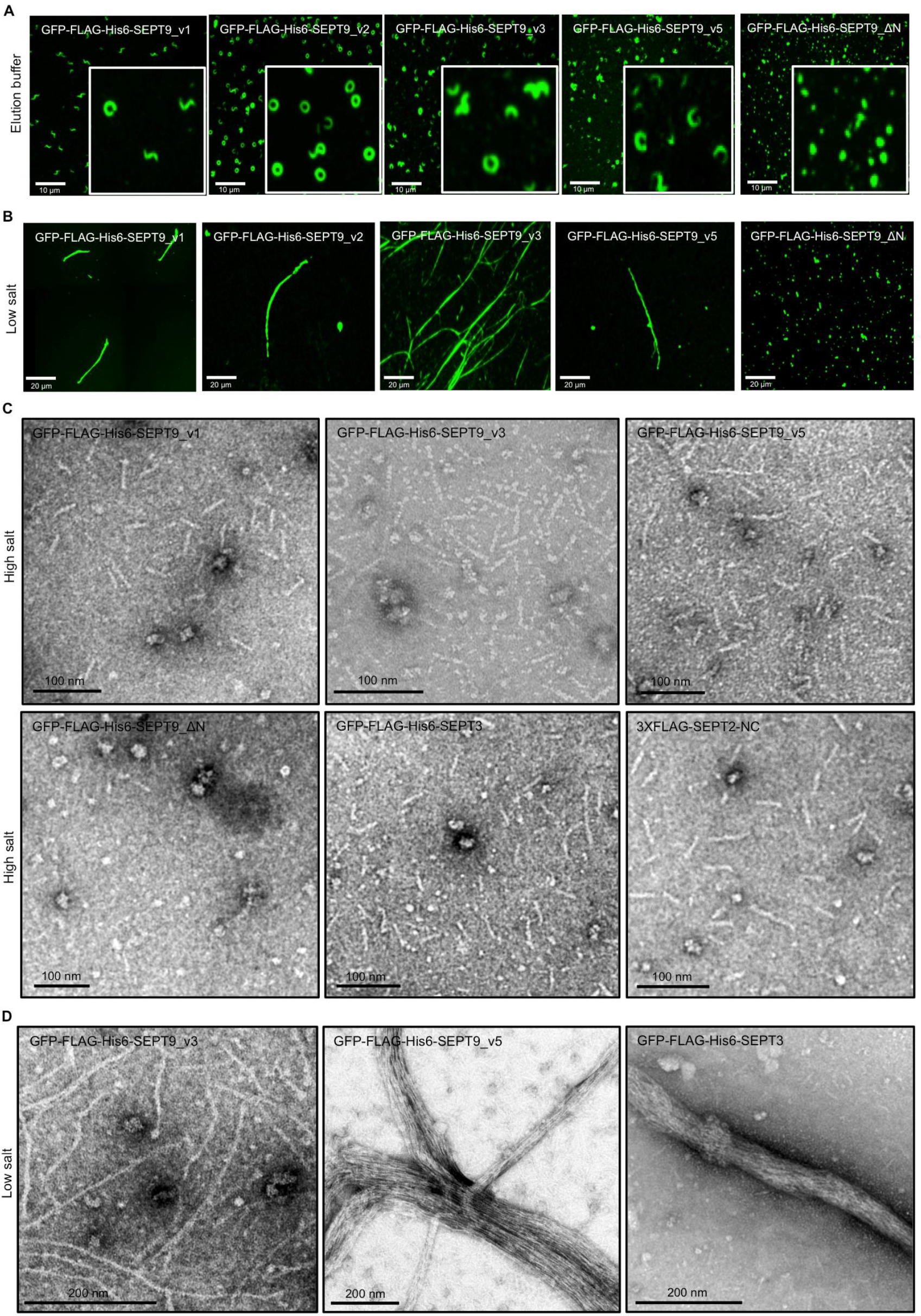
Octamers with both short and long isoforms of SEPT9 can polymerize in vitro but ΔN mutation blocks polymerization. (A) Fluorescence images of septin rings and curvilinear structures in vitro after purification from stable cell lines. 2μl of isolated complexes in elution buffer (150 mM KCl) spotted on the coverslips mounted and imaged. (B) Fluorescence images of septin filaments in vitro after dialysis of isolated complexes against low salt buffer (50mM KCl) for 24hrs. (C) Isolated complexes supplemented with high salt buffer (400mM KCl) and examined using negative staining EM. (D) Septin complexes isolated from inducible stable cell lines dialyzed against low salt buffer (50mM KCl) overnight followed by negative staining EM.

*In vitro* polymerization of these distinct septin complexes was assessed following dialysis into low salt buffer and imaging with electron and fluorescence microscopy. Our data suggest that septin complexes with different isoforms of SEPT9 polymerize to filaments *in vitro* (Figure 3B and 3D). Therefore, SEPT9_v4 and SEPT9_v5 with the shortest N-terminus do not seem to play a role as chain terminators. Interestingly, we found that SEPT9 isoforms with shorter N-termini (SEPT9_v4 or SEPT9_v5) were more frequently observed as bundled filaments (Figure 1F and 3D) while the longer isoforms were frequently single filaments.

Sellin and colleagues demonstrated that SEPT3 subgroup members (SEPT3, SEPT9, and SEPT12) are replaceable in the octamers (Sellin et al., 2014). We therefore set out to determine if SEPT3 containing complexes could polymerize *in vitro*. Inducible stable cell lines that express GFP-FLAG-His6-SEPT3 were generated and characterized by western blotting (Figure 2Aii). As expected the overexpression of SEPT3 caused a decrease in SEPT9 protein levels (Figure 2Aii). This corresponded with a significant decrease in SEPT9-containing octamers compared to uninduced cells (Figure 2Bii). Using siRNA targeting SEPT9, SEPT9 was further depleted in GFP-FLAG-His6-SEPT3 cells before doxycycline induction (Figure 2Aii). After isolating septin complexes from this cell line, we observed that they were indeed rod-shaped octamers and after performing an *in vitro* polymerization assay, they formed filaments as observed by EM (Figure 3D). Interestingly, we noticed that, like the short isoforms of SEPT9 (SEPT9_v4 and v5), SEPT3-containing complexes also favored bundle formation (Figure 3D).

We were unable to generate stable lines that expressed SEPT12, suggesting that this protein may be unstable in HeLa cells. Overall, while the extent of bundling may be different between isoforms, all tested SEPT3 subfamily members supported polymerization.

### Mutation analysis suggests that the canonical model of mammalian septin organization is inverted

In yeast, when Cdc11, which is located at the terminal position, is mutated at the N terminus, the octameric complexes fail to form filaments (Bertin and et al., 2008). If mammalian septin polymerization follows the same rules as yeast, and the SEPT3 subfamily occupies the terminal position, mutation of the N terminus should prevent this. Previously we reported that when SEPT9_ΔN mutants, which lack the N-terminus, were expressed in mammalian cells, septin filaments and higher order structures were disrupted (Kim et al., 2011).

To analyze how this mutation disrupts filament formation and/or complex assembly we generated inducible stable cell lines that express GFP-FLAG-His6-SEPT9_ΔN. Unlike the full-length SEPT9 variants, GFP-FLAG-His6-SEPT9_ΔN did not localize on actin stress fibers but were distributed in a punctate fashion throughout the cytoplasm (data not shown). After treating cells with the actin-disrupting drug cytochalasin D, GFP-FLAG-His6-SEPT9_ΔN did not incorporate into rings or curvilinear structures like their full-length counterparts (data not shown). Surprisingly, immunopurified GFP-FLAG-His6-SEPT9_ΔN complexes were tetramers under native conditions (Figure 2Bii). Complexes containing GFP-FLAG-His6-SEPT9_ΔN migrated well below the predicted hexamer band despite also containing SEPT2, SEPT6 and SEPT7. When spotted on coverslips in elution buffer, complexes isolated from GFP-FLAG-His6-SEPT9_ΔN cell lines were punctate in contrast to the rings, spiral and curvilinear structures seen for full length SEPT9 (Figure 3A). Moreover, when examined by EM in high salt buffer to prevent polymerization, the GFP-FLAG-His6-SEPT9_ΔN complexes were shorter than the rod-shaped octamers. After analysis and measurements of 500 of these structures, they fit the predicted length of tetramers with average lengths of <20nm (Figure S3). When dialyzed against the low salt buffer, GFP-Flag-His6-SEPT9_ΔN–containing tetramers failed to polymerize into filaments or higher order structures (Figure 3B and 3D). These unexpected results suggest that SEPT9 N-terminus removal prevents octameric complex assembly and are inconsistent with SEPT9 being located at the ends of octamers.

We therefore tested the effect of SEPT2 N terminal mutations on octamer structure. If SEPT2 was present in the middle of the octamer we would predict that its mutation should result in tetramer formation. We generated an inducible stable cell line that expresses 3X-FLAG-SEPT2-NC with a mutation at the NC interface (V27D) shown to prevent SEPT2-SEPT2 interactions (Sirajuddin et al., 2007; Kim et al., 2012). When septins from this cell line were resolved on blue native gels they revealed that 3xFLAG-Sept2-NC incorporated into complexes that are octamers and hexamers (Figure 2Biii) and of comparable length to wild type complexes by EM (Figure 3) but were unable to polymerize (data not shown).

Together, these results are most consistent with the notion that the order of septins in the octameric complexes is SEPT2-SEPT6-SEPT7-SEPT9-SEPT9-SEPT7-SEPT6-SEPT2.

### Mammalian septin hexamers and octamers associate *in vitro*

In the prevailing model of mammalian septin organization, hexamers would have the G-interface of SEPT7 exposed, while octamers would have the NC interface of SEPT9 exposed, and these should be incompatible for polymerization (Figure 4A). Our data proposes that hexamers and octamers would both have exposed SEPT2 NC interfaces with SEPT9 missing from the middle of the complex and would therefore be predicted to co-polymerize. This is analogous to the situation in *S. cerevisiae*, where Cdc10 (the yeast septin most like SEPT9) is present in the middle of the complex, and yeast lacking Cdc10 can form hexameric complexes that retain the ability to polymerize and support viability (McMurray et al., 2011). To test hexamer-octamer co-polymerization we generated mCherry-FLAG-SEPT7 cell line to isolate red hexamers after depleting endogenous SEPT9 (Figure 4Bi, 4Bii, and 4Biii). Like the GFP versions, purified complexes containing mCherry-FLAG-SEPT7 are capable of polymerization alone (Fig 4Biii) and when mixed with GFP-FLAG-His-SEPT9_v3 resulted in the formation of mixed filaments that appeared yellow in merged images (Figure 4C) indicating nearly complete co-localization (Pearson coefficient correlation: 0.75, p<0.01). This suggests that mammalian septin hexamers and octamers can co-polymerize in vitro, consistent with our new model.

**Figure 4:**
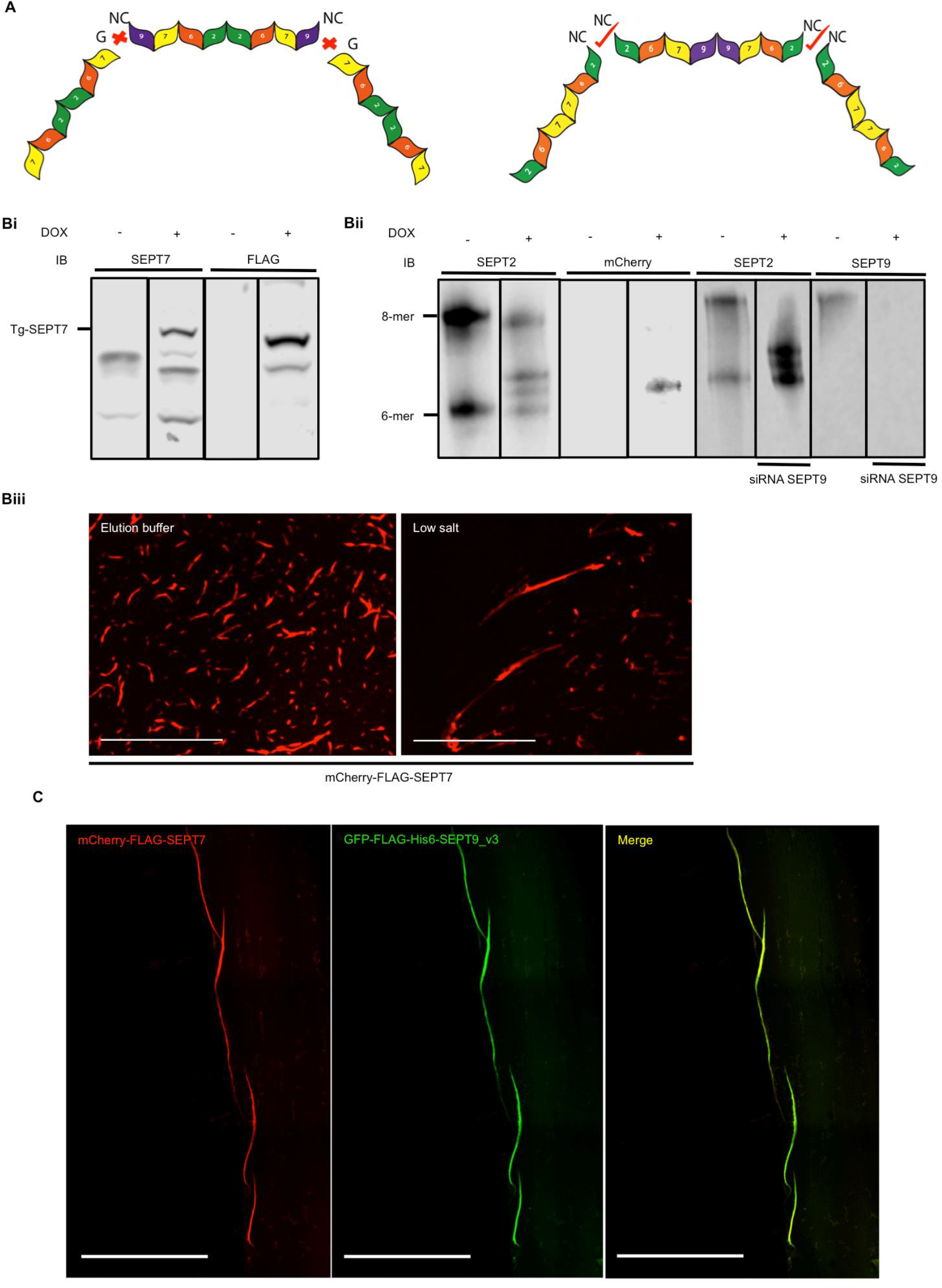
Mammalian septin hexamers and octamers associate in-vitro. (A) Schematic representation of the former model of hexameric and octameric human septin complex suggest incompatible ends unable to interact (left) and an alternative arrangement where compatible ends would support co-polymerization. (Bi) western blot of induced mCherry-FLAG-SEPT7 cell lysate immunoblotted for SEPT7 and FLAG antibodies. (Bii) Blue native PAGE of induced and uninduced mCherry-FLAG-SEPT7 cell lysates resolved by blue native PAGE followed by immunodetection with SEPT2 and mCherry antibodies. In the induced lane there are three bands detected with SEPT2 antibodies. The lower band, which also appears in the uninduced lysate, is the endogenous hexameric band. mCherry-FLAG-SEPT7-containing hexamers are shifted and were also detected by the mCherry antibody. Depletion of SEPT9 eliminates octamers but had no effect on hexamers. (Biii) Isolated complexes from mCherry-FLAG-SEPT7 inducible stable cell line in the elution buffer (150mM KCl) and low salt buffer (50mM KCl) were spotted on the coverslips and examined with fluorescence microscopy. (C) Green octamers and red hexamers were mixed after dialysis with low salt buffer (50 mM KCl) and examined with fluorescence microscopy. Green and red channels show near perfect co-localization as indicated by Yellow in the Merge.

### Borgs influence septin polymerization

Borgs (Binder of Rho GTPases) (also known as CDC42EP1-5) are Cdc42 effector proteins that were previously shown to bind and regulate septins. Overexpression of Borgs appeared to change the higher-order structure of septins by promoting thicker bundled filaments. Cdc42-GTP binding to Borgs, however, has been shown to negatively regulate this interaction (Joberty et al., 1999, 2001). Gic1 in yeast appears to be a functional homolog of the human Borg proteins and is also a Cdc42 effector. When Gic1 is added to purified yeast septins (Cdc3-EGFP, Cdc10, Cdc11, and Cdc12), it enhances their polymerization and leads to cross-linking of filaments (Sadian et al., 2013). Interestingly, when Cdc42 was added to Gic-septin filaments, dissociation of septins from Gic1 was observed. This suggests that, in agreement with the previous results for Borgs in mammalian cells, Cdc42 negatively regulates the interaction of Gic1-septins in yeast (Joberty et al., 2001; Sadian et al., 2013). In the same study, it was determined that the binding site of Borgs on the yeast septin complex is at the innermost septin Cdc10, and the lateral spacing between filaments cross-linked by Gic1 was measured to be 20nm. To date, no *in vitro* assay has examined how Borgs alter the structure of the mammalian septin complex.

Borgs bind to septins via the Borg homology domain 3 (BD3). The BD3 domain, located at the C-terminus of Borg3, was shown to be sufficient to bind septins and mutation of a hydrophobic cluster of amino acids (L102A, V105A, L106A) blocks this interaction (Joberty et al., 2001). Additionally, it was shown that Borgs bind to the C-terminal coiled-coil domains of SEPT6 and SEPT7 (Sheffield et al., 2003).

To begin to analyze the effect of Borgs on mammalian septin polymerization *in vitro*, we conducted an *in vitro* polymerization assay using isolated hexamers from mCherry-FLAG-SEPT7 stable cell lines depleted of SEPT9. In the absence of the BD3 domain, consistent with previous observations, the hexamers form single filaments. In the presence of BD3, the structure of polymerized filaments was altered, and train track-like filaments with cross-bridges were observed (Figure 5A right panels). This structure was very similar to that observed when Gic1 was added to yeast septin complexes. We also frequently observed ring-like structures by EM (Figure 5A) although the significance of this is not known. In contrast, addition of BD3 to octameric complexes containing GFP-FLAG-His6-SEPT9_v3 or GFP-FLAG-His6-SEPT9_v5 resulted in septin filaments bundled into cable-like structures with multiple filaments per bundle appearing to surround pronounced cross bridge densities much larger than seen for the hexamers (Figure 5A left panels and S4). To further confirm that the BD3 domain is responsible for changing septin polymerization, we performed the same experiment in the presence of recombinant BD3 where the LVL residues were mutated to alanines. As expected, this mutant could not promote the formation of cable-like filaments with cross bridging (Figure 5C).

**Figure 5:**
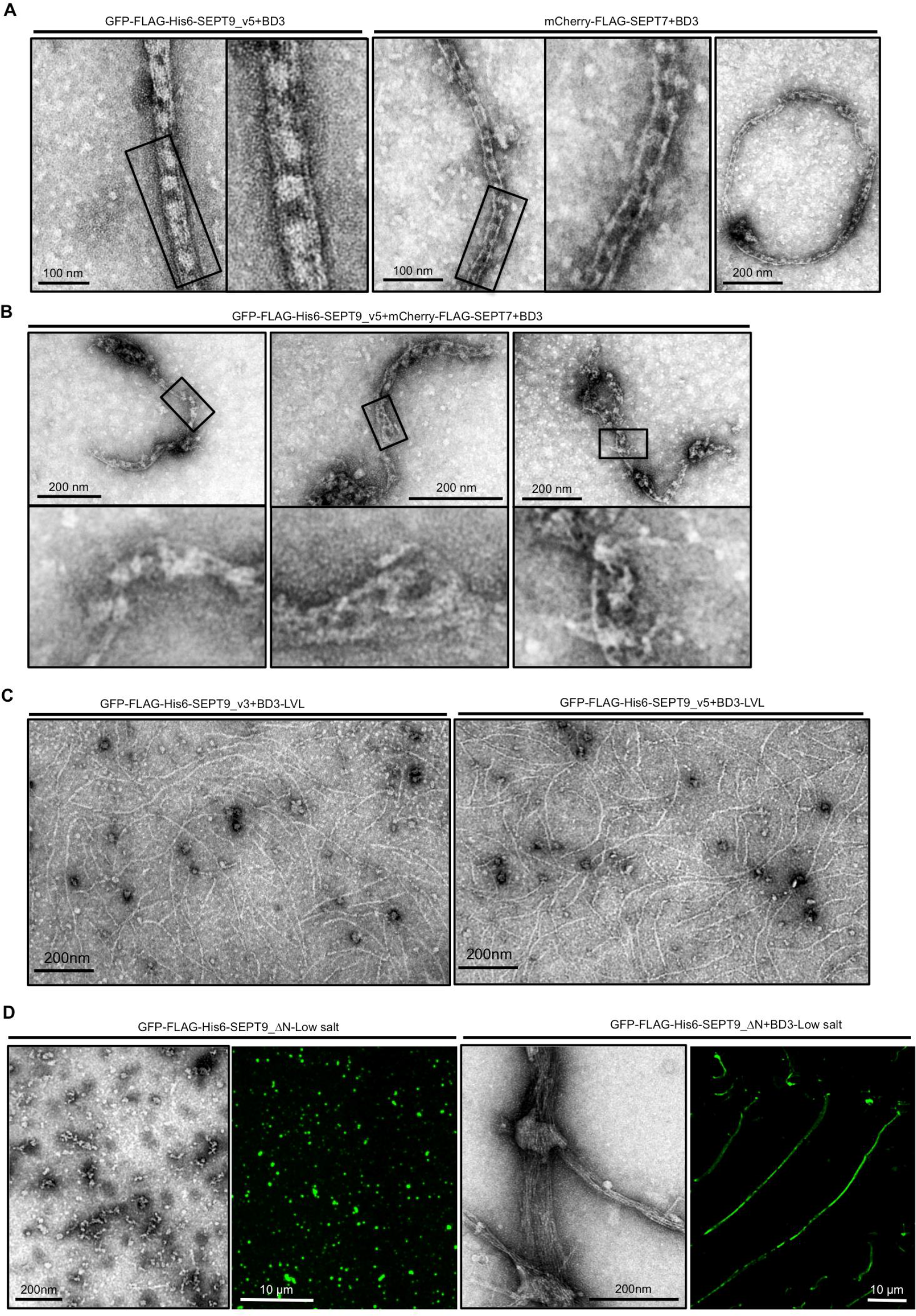
BD3 domain alters the polymerization of mammalian septin hexamers in-vitro. (A) BD3 differentially bundles hexamers and octamers. BD3 domain was added to the mCherry-FLAG-SEPT7 or GFP-FLAG-His6-SEPT9_v5-containing complexes and dialyzed against low salt buffer (50mM) followed by negative staining EM. Cable-like structures and rings were observed in the presence of the BD3 domain, but more prominent cross-bridges and more filaments appear present in octameric filaments. (B) BD3 addition to mixed filaments formed from mCherry-FLAG-SEPT7 and GFP-FLAG-His6-SEPT9_v5 were irregular in structure, and often containing loops, flares and frays, unlike the regular structures seen in A. (C) BD3-LVL mutants did not promote septin cable-like structures. (D) BD3 promotes polymerization of GFP-Flag-His6-SEPT9_ΔN-containing complexes as demonstrated by fluorescence microscopy and negative staining EM.

To determine the periodicity of the cross-bridges, we applied a fast fourier transform (FFT) algorithm to density scans along the length of the filaments. Interestingly, the periodicity of the cross bridging in octamers was approximately 35nm (Figure S4). This result was surprising since if BD3 binds to the coiled-coil domains of SEPT6 and SEPT7, this distance would be predicted to be shorter (Figure S4).

It is interesting to note that the cross-bridges are more robust for octamer bundles than for hexamers. In yeast, the cross-bridges were composed of Gic1 protein copies, and since the octamers and hexamers bundled differently in the presence of BD3, this raised the possibility that, in addition to the SEPT6 and SEPT7 coiled coils, BD3 may bind to SEPT9. If so, Borgs might bind to two adjacent sites, at the SEPT6/7 coiled coils, and to SEPT9, giving rise to a large protein mass that would have a centroid around the SEPY9-SEPT9 interface and a periodicity of 35 nm. To elucidate whether SEPT9 binds to Borgs, FLAG-His-tagged versions of SEPT2, SEPT6, SEPT7, SEPT9, or SEPT6+SEPT7 were transiently co-transfected with GFP-Borg3 in HeLa cells. Following His pull down, western blot analysis confirmed that FLAG-His-SEPT6 and FLAG-His-SEPT6+SEPT7 pulled down GFP-Borg3 but FLAG-His-SEPT9 did not (Figure S4). Since the presence of SEPT9 clearly changes the BD3 binding behavior, this result was unexpected and we, therefore, considered the possibility that BD3 may bind at the interface between SEPT7 and SEPT9. Hence, we co-expressed FLAG-His-SEPT7+SEPT9 and GFP-Borg3 (Figure S4). However, even the SEPT7/SEPT9 heterodimer could not pull down GFP-Borg3 (Figure S4). Unfortunately, we were unable to demonstrate direct binding of SEPT9 binds to Borgs using the pull-down assay.

We have observed that the cross-bridges are periodic when either hexamers or octamers are polymerized in the presence of BD3 but appear different from each other. However, if hexamers and octamers interchangeably and randomly co-polymerize within single filaments, then we would not expect to see regular periodicity when BD3 bundles mixed filaments since filament registry would be lost. Indeed, as shown in Figure 5B, although cable-like filaments do form, in most of these structures there are regions where the periodicity is largely lost, and the bundles have irregular loops, frays and disorganized tangles. These results further support the likelihood that hexamers and octamers co-polymerize and show that their bundling in the presence of BD3 is much more complex than for filaments of pure hexamers or octamers where lateral registry can be achieved.

As discussed above, GFP-FLAG-His6-SEPT9_ΔN-containing complexes are not able to form filaments *in vitro* and they are tetrameric (Figure 3B, 3C, and 2Bii). Similar ΔN mutations have been used in many studies and shown to prevent filament formation. Therefore, it was surprising to see that adding BD3 to GFP-FLAG-His6-SEPT9_ΔN-containing complexes led to their polymerization *in vitro* (Figure 5D). Interestingly, these bundled filaments lack the cross-bridge structures seen in octameric complexes containing full length SEPT9.

## DISCUSSION

In this study we have utilized purified mammalian hexameric and octameric septin complexes to test the role of SEPT3 family members in septin filament polymerization, but these studies led to surprising revelations about the organization of septins within the hetero-oligomeric complexes.

### Septin hexamers and octamers can polymerize *in vitro*

Previous studies, confirmed herein, revealed that mammalian cell lines express both hexamers and octamers (Sellin et al., 2011). We had previously shown that depletion of SEPT9, which would shift the balance to only hexamers, resulted in abscission defects in late telophase, whereas depletion of core hexamer/octamer septins (for example SEPT2) resulted in defects in the stability of the actomyosin contractile ring at earlier stages of division (Estey et al., 2010). In contrast, overexpression of SEPT9 can change the balance in favor of octamers over hexamers, and overexpression of specific isoforms of SEPT9 has been associated with sporadic breast and ovarian cancers (Gonzalez et al., 2007; Scott et al., 2006), supporting the distinct roles of these complexes. Here we show that both hexameric and octameric complexes purified from mammalian cells can polymerize under conditions of low salt into long, linear filaments, implying that functional distinctions do not lie in their abilities to polymerize, but may reflect their ability to interact with other proteins or signaling complexes. Octamers comprised of different isoforms of SEPT9, or subfamily member SEPT3, were all able to polymerize, although those with shorter N-termini (SEPT9_v4, _v5, SEPT3) appeared to bundle laterally to a greater extent. We previously showed that SEPT9_v4 could not rescue the cytokinesis defect caused by general SEPT9 knockdown (Estey et al., 2010). This could indicate that the longer isoforms of SEPT9 interact with proteins critical for this process, or perhaps that these thick bundles create septin structures that are not suitable to complete cytokinesis.

### The canonical model of mammalian septin organization is inverted

The canonical model of mammalian septin organization based on the crystal structure of hexamers (Sirajuddin et al., 2007) placed SEPT2 in the middle of both octamers and hexamers, while SEPT7 was at the ends of hexamers exposing a G-interface while SEPT9 would be at the ends of octamers exposing an NC interface. This implied that hexamers and octamers should not be able to interact and that two forms of filaments would exist in mammalian cells. Surprisingly, mutation of the SEPT9 interaction surface resulted in tetramers that could not polymerize (Figure 2Bii and 3B), while depletion of SEPT9 by siRNA resulted in hexamers (Figure 4Bi). Intriguingly, mutations in the NC interface of Cdc10, which prevents Cdc10-Cdc10 homodimer formation, causes tetramer formation and loss of viability, while null Cdc10 strains form hexameric complexes that support viability (McMurray and et al., 2011). Cdc10 is the yeast septin most structurally similar to SEPT9. In contrast, mutation of the SEPT2 NC interface gives rise to hexamers and octamers that cannot polymerize (Figure 2Biii and Figure 3C and data not shown). Similarly, mutation of the NC interface of the terminal Cdc11 subunits results in octameric complexes that cannot polymerize and do not support survival (McMurray et al., 2011).

Hence, this study upends the existing model and proposes an alternative conformation for mammalian septin octamers and hexamers arranged as SEPT2-SEPT6-SEPT7-SEPT9-SEPT9-SEPT7-SEPT6-SEPT2 and SEPT2-SEPT6-SEPT7-SEPT7-SEPT6-SEPT2 respectively. This new model is not only consistent with the currently known yeast complex but would provide compatible ends that would permit hexamers and octamers to co-polymerize, as we observed. It would also fit with the stepwise model of septin polymerization proposed by Weems and McMurray (Weems and McMurray, 2017). Briefly, they propose that Cdc3, Cdc12 and Cdc11 form trimeric complexes that can then assemble with a Cdc10-Cdc10 dimer to form an octamer. By analogy, we speculate that SEPT9 would form a dimer at the core of mammalian septin complex that then interacts with the 7-6-2 trimetric forms to form the octameric complex arranged as 2-6-7-9-9-7-6-2. In this scenario when SEPT9 is absent the two trimetric halves come together to form hexameric mammalian septin complex. It should be noted that the trimeric 2-6-7 and dimeric 9-9 complexes are not detected on blue native gels and therefore must exist only very transiently. In order to enhance the correct dimerization at the G-interface of SEPT7-SEPT7 in hexamers, and perhaps to control polymerization, there must be a mechanism to prevent NC-interactions of SEPT2 prior to hexamer/octamer formation. Interestingly, our lab has previously shown that mutation of a single amino acid in the N terminus of SEPT2 causes excessive homotypic polymerization of overexpressed SEPT2 in cells (Kim et al., 2012), suggesting that it may have a role in inhibiting SEPT2-SEPT2 interactions under normal conditions.

The crystal structure of the hexameric septin complex was solved after expressing SEPT2, SEPT6, and SEPT7 in a bacterial expression system. The crystal contained repeats of alternating trimers and the order of the hexameric complex was based on class averages of negative stain EM images where SEPT2 was tagged with MBP. However, the densities of MBPs were not clearly defined in these images and it is possible that the authors incorrectly identified their positions. On the other hand, it has been shown that in the absence of preferred septin partners, septins can bind at non-conventional interfaces. For instance, in the hexameric complex SEPT2-SEPT2 interaction is via their NC interface but SEPT2-SEPT2 dimers interact via their G interfaces when expressed alone (Sirajuddin et al., 2007). Since SEPT9, which is the preferred partner of SEPT7, was not expressed in bacteria, it is possible that the two trimeric halves came together in an unconventional way with SEPT7 at the ends and therefore presented a model of an artifactual complex. Since our complexes are isolated from mammalian cells we feel confident that the organization represents the situation *in vivo*.

### BD3 domain is sufficient for cable-like structure formation in vitro

The BD3 domain of Borgs has been shown to bind and bundle septin filaments in cells (Joberty et al., 2001), but how it does so was not known. Here we show a dramatic change in the structure and appearance of septin filaments when BD3 was present, but not if the septin-binding LVL motif was mutated to alanine. Moreover, the bundled appearance was different for filaments comprised of hexamers and octamers. The former appeared as train-track pairs of filaments while the latter resembled cable–like bundles. In both cases cross-bridges, presumably comprised of BD3, appeared at regular intervals but were much larger for the octamers. This is strikingly reminiscent of the bundled filaments seen for yeast septins when mixed with Gic1 (Sadian et al., 2013). 3-D reconstruction of yeast septin filaments in the presence of Gic1, which is a Borg homolog, demonstrated that approximately 6 filaments and 12 Cdc10 are involved. Also, Gic1 elution from gel filtration has shown that Gic1 forms dimers (Sadian et al., 2013). Therefore, a detected cross-bridge might include 12 Gic1 dimers forming a dense mass. Notably, this finding was contradictory to a previous report that Gic1 binds to Cdc12 (Iwase et al., 2006), raising the possibility of more than one binding site.

In pull downs BD3 can bind to the coiled-coil domain of SEPT6 and SEPT7 (Figure S4 and Sheffield et al., 2003). However, the fact that octamers interact differently with BD3 than hexamers suggests that BD3 may also interact with SEPT9, although we were unable to demonstrate this. In the case of octamers, these cross-bridges were spaced at about 35 nm apart, consistent with the length of octameric subunits but inconsistent with the known binding site of BD3 to the coiled-coils of SEPT6 and SEPT7. We speculate that perhaps undefined binding sites on SEPT9 may lead to a cluster of BD3 molecules at SEPT9 and SEPTs 6+7 that appear as a single mass in negative staining, once per repeat, leading to a spacing that is similar to the length of the octamer (Figure S4C).

Surprisingly, BD3 was able to rescue the polymerization of GFP-Flag-His6-SEPT9_ΔN, yet they formed filaments without cross-bridges. We can only speculate as to how BD3 supported polymerization yet did not appear to incorporate into bundled structures. For example, it is possible that BD3 binding stabilized the interactions between tetramers, allowing a longer “window of opportunity” for their assembly. This would be analogous to the model proposed by Schaefer and colleagues where an extended G1 cycle stage allowed mutant forms of the septins to be incorporated into functional octamers that would not otherwise assemble under normal folding kinetics (Schaefer et al., 2016). Alternatively, BD3 domain dimers may be forming a bridge between tetramers to create pseudo-octamers that can then polymerize like normal octamers. Since BD3 was shown to bind to the coiled-coil complex between SEPT6 and SEPT7, this should have still occurred in the tetramers, but would have been expected to produce aggregates or clusters, which we did not observe.

Further work will be required to identify the structural details of Borgs and septins and to elucidate the number of BD3 proteins and septin filaments involved in the cable-like filaments.

## MATERIALS AND METHODS

### Cell Culture and Drug Treatment

Flip-in HeLa cells stably expressing GFP-FLAG-His tagged septins and wild-type was generated and subcultured with 5% FBS and Dulbecco’s Modified Eagle’s Medium (DMEM) in 37°C in a humidified incubator with 5% CO_2_. Cells seeded onto two-175cm2 tissue culture flasks were grown to ~50%-60% confluence the next day. Addition of 100ng/mL doxycycline induced the expression of the transgene. After 24hrs cells were trypsinized, pelleted and washed twice in PBS. The pellet then flashed frozen and kept at −80°C prior to purification.

For the disruption of actin filaments, induced cells were treated with 3 μM cytochalasin D (Sigma-Aldrich) 1hr prior to starting the Immunofluorescence experiment.

### Constructs

GFP-FLAG-His6-SEPT9 variants were cloned into a modified inducible mammalian FRT vector (pcDNA5 FRT-TO). The hygromycin cassette was replaced with puromycin, for rapid generation of stable cell lines. All SEPT9 variants were cloned using Gibson Assembly cloning (NEB#E5510S) according to manufacturer’s recommendations. Briefly, septins were amplified with PCR, and gel extracted. The amounts of insert and vector were adjusted accordingly to ensure a molar ratio of 1:1. The DNA fragments and the Gibson Assembly Master Mix (2X) was combined in a total volume of 4μL and incubated at 50°C for 30 minutes. The entire reaction was transformed into competent bacteria (DH5α) and plated on ampicillin LB-agar. All constructs were sequenced in both directions. The accession numbers of all five SEPT9 variants are: NM_001113491, NM_001113493, NM_006640, NM_001113492 and NM_001113496. Certain isoforms (V1, V3 and V4) were described previously (Estey et al., 2010, Kim et al., 2011). SEPT9_V2 was generated using SEPT9_V3 as template and an extended forward primer (5’-atatGAATTCatgtcggaccccgcggtcaacgcgcagctggatgggatcatttcggacttcgaagCCTTGAAAAGATCTTTTGAGGTCG-3’). Since SEPT9_V5 can be thought of as an N-terminal truncation of SEPT9_V4, it was amplified from a V4 template using appropriate primers.

### Blue native PAGE

Cell lysates were resolved using blue native PAGE following the protocol as described (Sellin et al., 2014), that ensures the complete disassembly of septins to their core complexes. Cells were grown in a T75 flask and induced with 100ng DOX for 24 hrs or not induced, then trypsinized and washed with PBS. The permeabilization buffer (0.2% saponin, NaCl 0.45 M, 80 mM PIPES (pH6.8), 5 mM MgCl2, 4 mM ethylene glycol tetraacetic acid (EGTA), 0.1mM guanosine 5’-triphosphate (GTP), 25 mM ε-aminocaproic acid, and protease inhibitors) was added to the cell pellet to yield a final concentration of 3 to 6 mg/ml soluble protein. Cells were passed through a syringe (G71/2) 3X and centrifuged at 14,000 X g for 30 min. During the centrifugation, the desalting columns (30 kDa cut-off; Micro Bio-Spin P30 Column; Bio-Rad) were equilibrated with desalting buffer (40 mM BisTris-HCl pH 7.2, 0.3 M NaCl, 1mM ethylene glycol tetraacetic acid (EGTA), 25 mM ε-aminocaproic acid, 0.2mM MgCl2). The supernatants from the last centrifugation step were passed over a desalting column. A bicinchoninic acid assay was performed to determine the concentration of soluble protein in each sample. Cell extracts were diluted in dilution buffer (50 mM BisTris-HCl, 50 mM NaCl,6% sucrose, 0.05% Coomassie Blue G250, 250 mM aminocapric acid) to 0.4 μg/μl and 20 μl was loaded per well. For resolving native septin complexes 4–16% Novex NativePAGE Bis-Tris Gel System precast poly¬acrylamide gels (Invitrogen, Carlsbad, CA) were used. The electrophoresis was done based on the protocol provided by the manufacturer using Native page running buffer (20x) (Bis-Tris 50mM, Tricine 50mM), Native page cathode additive (20x) (0.4% Coomassie G-250) to the final concentration of 1X. For western blot, proteins were transferred to PVDF membranes overnight at 100 MA using a non-denaturing transfer buffer (31 mM Tris-HCl, 300 mM Glycin). The rest of the western blot procedure was done according to the same protocol described in the previous section.

### Purification of Mammalian Flag-Septin Complexes

Our protocol for isolating mammalian septin heteromers is adapted from “Sigma Product Technical Bulletin” and a protocol by Sellin et al. (Sellin et al., 2011). Pre-cooled Permeabilisation Buffer (PEM (80mM PIPES pH6.9, 4mM EGTA, 2mM MgCl2, 150 mM KCl) with 0.2% saponin), was added to the cell pellet and mixed by pipetting up and down and rotating in a 4°C for 30 min. After centrifugation at 500 Xg for 5 min the soluble fraction was collected in a new falcon tube and supplemented with 20% Triton, centrifuged at 12000 Xg for 10 min cold and the supernatant was transferred to the fresh falcon tube. During the centrifugation, 40µL of slurry protein G agarose beads (Sigma) washed 2X with 1X TBS (50mM Tris-HCl pH7.4, 150mM NaCl) and 1X Lysis buffer at 1000g for 30sec. To pre-clear the supernatant from the last step, they were mixed with washed protein G agarose beads and incubated 10 min at 4°C on a nutator. After centrifugation at 1000 Xg for 1 min, pre-clear supernatant was transferred to the fresh 10ml falcon tube. 50 µL of Flag-agarose gel slurry (Anti-Flag M2 Affinity Gel (Sigma, Cat#: A2220)) was transferred to a 1.5mL microfuge tube washed 2X with 1XTBS and 1X Lysis buffer at 1000 Xg for 30sec. The cleared supernatant from the last step was then incubated with FLAG beads at 4°C on a nutator for 2hrs.

The mixture of FLAG beads and pre-cleared lysate was centrifuged at 1000 Xg for 1 min and the supernatant was carefully removed. The FLAG beads were then washed extensively with pre-cooled wash buffer (PEM with 0.1% Triton X-100, 150 mM KCl) at 4°C. FLAG beads were then incubated with Elution buffer PEM with an additional 4mM EGTA, 1mM GTP, a protease inhibitor tablet, 150 mM KCl and 100ug/mL FLAG peptide) at RT for 15 min and periodically tapping with a finger. The mixture was then Spin down 1000 Xg for 2 min and the eluate was carefully removed with a 27-gauge needle and transferred to a fresh falcon tube.

### Electroporation

SEPT9 siRNA that targets this sequence 5’-GCACGATATTGAGGAGAAA-3’, previously shown to effectively deplete Septin9 (Estey et al., 2010) (Dharmacon) was dissolving in diethyl pyrocarbonate (DEPC)-treated water at a concentration of 40 pmol/µL, stored at −80°C.

For depletion of endogenous SEPT9 in the mCh-FLAG-SEPT7 cell line, SEPT9 siRNA was electroporated into the cells. After 48hrs cells were induced with 100 ng of DOX and after 72 hrs cells were frozen prior to protein purification. SEPT9 depletion was confirmed by immunoblotting with an antibody against SEPT9

### Immunoblotting

Immunoblotting was performed following standard protocols with 1hr of blocking, primary and secondary antibody incubations. The following primary antibodies were used: rabbit anti-SEPT9 (Proteintech) at 1:5000, rabbit anti-SEPT7 (Proteintech) at 1:5000, mouse anti-SEPT2 (Proteintech) at 1:7500, rabbit anti-SEPT6 (Proteintech) at 1:5000, rabbit anti-GFP at 1:5000, rabbit anti-SEPT11 (generated in house) at 1:1000, mouse anti-FLAG at 1:7,500 (M2; Sigma-Aldrich). Secondary, HRP-conjugated antibody was used at 1:5000.

### Immunofluorescence Staining

After washing with PBS, cells were fixed in 4% paraformaldehyde in PBS for 20 min at RT, then inactivated and permeabilized with 50mM NH4, 50mM glycine, and 0.1% TritonX-100 in PBS pH 7.4 for 20min at RT. After blocking cells with 5% horse serum in PBS primary (Rabbit anti-GFP at 1:200) and secondary antibodies were added for 1hr incubations with wash steps between. Coverslips were then mounted with DAKO fluorescent mounting medium (DAKO Canada) and allowed to dry overnight at RT in the dark.

Slides were imaged on an Olympus IX81 Quorum Spinning Disk Confocal microscope at The Hospital for Sick Children’s Imaging Facility, Toronto, ON. Images were acquired with Volocity software (Perkin Elmer) using a 60 X oil immersion objective.

### Electron microscopy

Isolated septin complexes were supplemented with high salt buffer (1X PEM, 400Mm KCl) to ensure all septin higher order structures are disassembled to septin complexes. For generating BD3-septin complexes, BD3 was added to the in vitro polymerization assay based on the method described by Sadian (Sadian et al., 2013).

Samples were then adsorbed on the glow-discharged carbon-coated copper grids for 30 seconds followed by three sequential washes (DDH_2_O) and staining with either 2% uranyl formate or 2% uranyl acetate.

For imaging septin filaments and higher order structures, isolated complexes were dialyzed against low salt buffer (50mM KCl) overnight and the rest of procedure was as described above. Samples were imaged at The Hospital for Sick Children’s Nanoscale Biomedical Imaging Facility using an electron microscope (TECNAI T20; FEI) operated at 200 kV. Length measurements of rods were performed using ImageJ

## ACKNOWLEDGEMENTS

This work was supported by grants from the Natural Sciences and Engineering Research Council and the Canadian Institutes of Health Research (MOP133660) to WST. WST is the recipient of the Canada Research Chair in Molecular Cell Biology. FS was supported by a RESTRACOMP award from the Hospital for Sick Children.

---

**Supplementary Figure 1:**
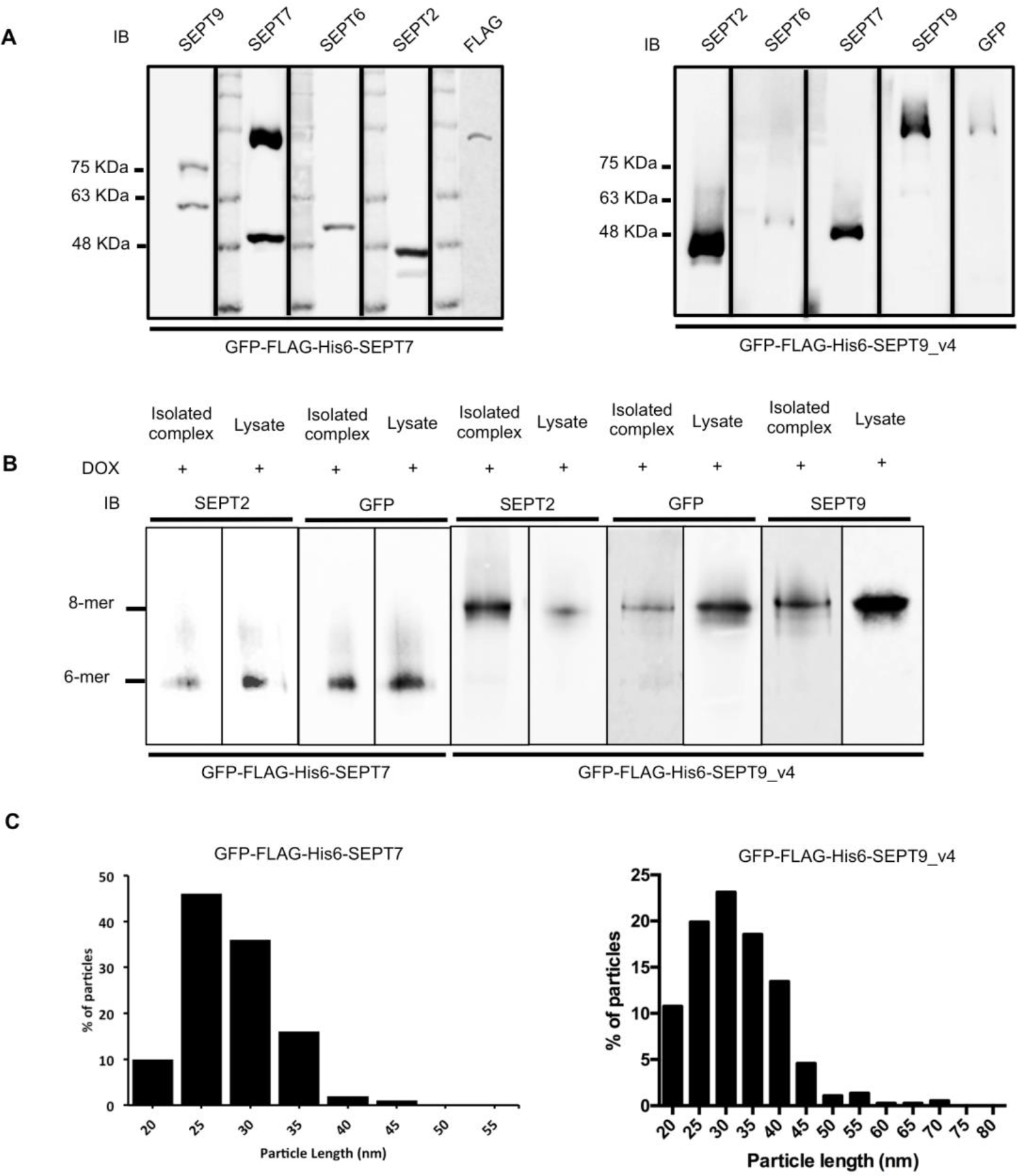
(A) Isolated complexes from GFP-Flag-His6-SEPT7 and GFP-Flag-His6-SEPT9_v4 immunoblotted with FLAG or GFP, SEPT2, SEPT6, SEPT7, and SEPT9 antibodies to confirm the location of septin bands in the Coomassie blue–stained SDS–PAGE gel. (B) Isolated septin complexes were run on native PAGE gel adjacent to the cell lysate and immunoblotted for SEPT2, SEPT7, SEPT9, and GFP to determine that isolated complexes were hexamers octamers. (C) Quantification of septin complexes isolated from GFP-FLAG-His6-SEPT7 and GFP-FLAG-His6-SEPT9_v4 stable cell lines in high salt buffer (400mM KCl) after negative staining EM.

**Supplementary Figure 2:**
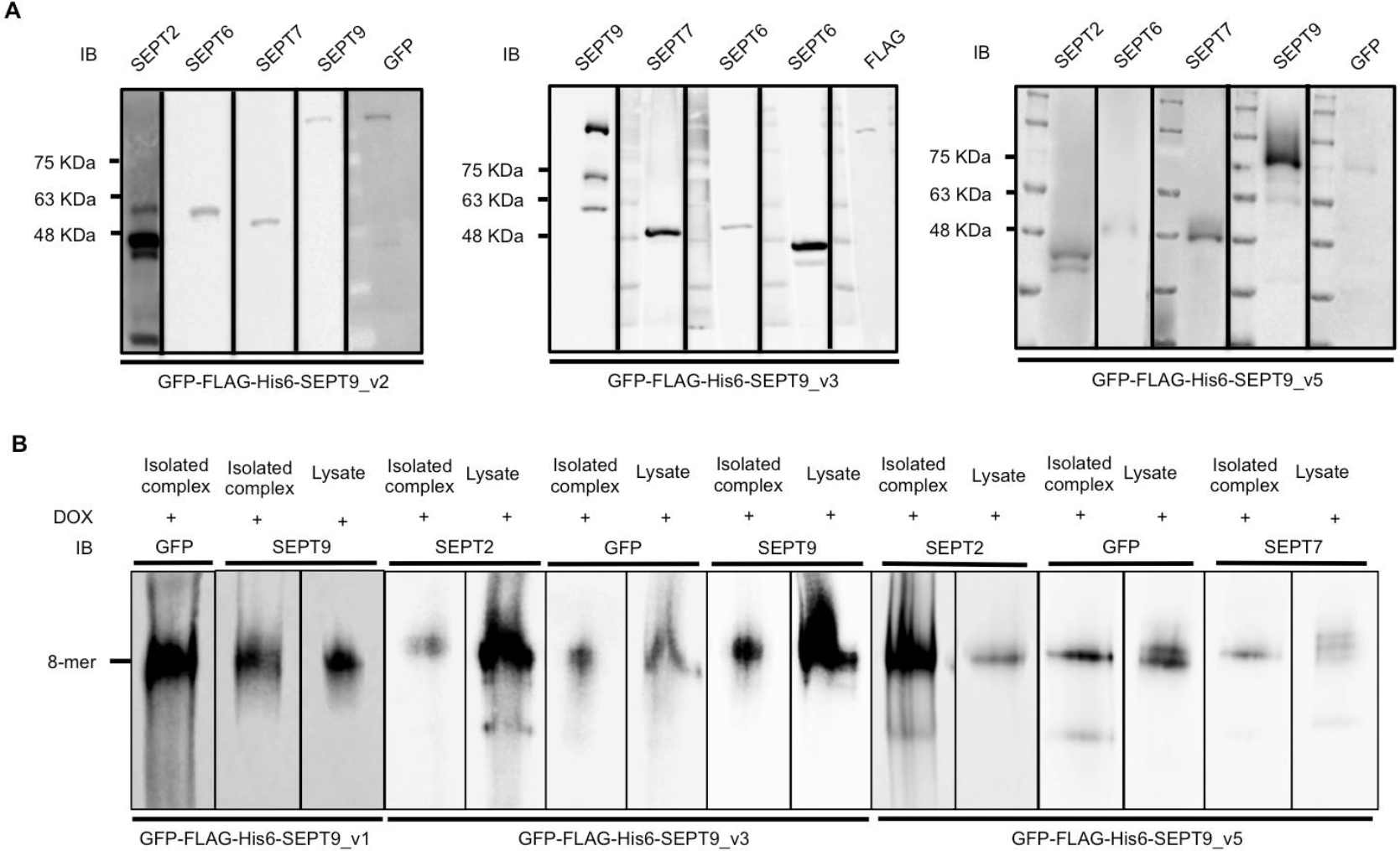
(A) Immunofluorescent staining of GFP (green) and actin (red) and colocalization of GFP with phalloidin-stained actin stress fibers (merge). The white-boxed areas are magnified images of expressed SEPT9 and actin stress fibers. (B) Isolated complexes from GFP-Flag-His6-SEPT_v1, GFP-Flag-His6-SEPT9_v2, GFP-Flag-His6-SEPT9_v5 immunoblotted with FLAG or GFP, SEPT2, SEPT6, SEPT7, and SEPT9 antibodies to confirm the location of septin bands in the Coomassie blue–stained SDS–PAGE gel. (C) Isolated septin complexes from GFP-Flag-His6-SEPT_v1, GFP-Flag-His6-SEPT_v3, and GFP-Flag-His6-SEPT_v5 were run on native PAGE gel adjacent to the cell lysate and immunoblotted for SEPT2, SEPT7, SEPT9, and GFP.

**Supplementary Figure 3:**
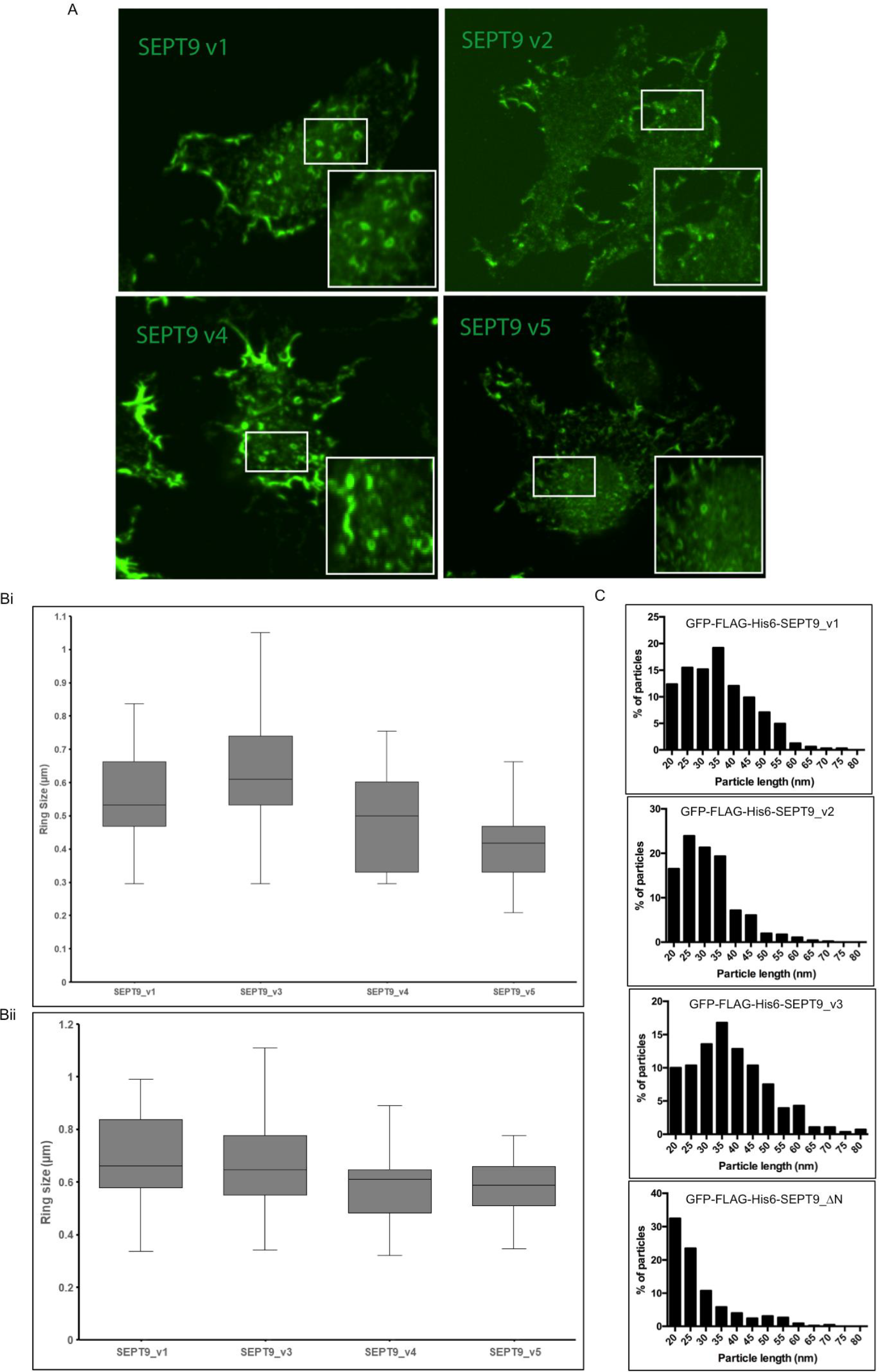
(A) Immunofluorescence staining of GFP (green) after disruption of actin-templated septin fibers upon cytochalasin D treatment. Septin rings magnified in boxed areas. (Bi) Quantification of septin rings in GFP-FLAG-His6-SEPT9_v1, GFP-FLAG-His6-SEPT9_v3, GFP-FLAG-His6-SEPT9_v4, and GFP-FLAG-His6-SEPT9_v5 stable cell lines after CD treatment. (Bii) Quantification of septin rings in GFP-FLAG-His6-SEPT9_v1, GFP-FLAG-His6-SEPT9_v3, GFP-FLAG-His6-SEPT9_v4, and GFP-FLAG-His6-SEPT9_v5 stable cell lines after isolating septin complex from these cell lines. (C) Isolated complexes from GFP-FLAG-His6-SEPT9_v1, GFP-FLAG-His6-SEPT9_v2, GFP-FLAG-His6-SEPT9_v3, and GFP-FLAG-His6-SEPT9_ΔN supplemented with high salt buffer (400 mM KCl) followed by negative staining EM. Quantification of septin complexes shows that isolated complexes from GFP-FLAG-His6-SEPT9_ through _v3 were octamers whereas those of GFP-FLAG-His6-SEPT9_ΔN were tetramers.

**Supplementary Figure 4:**
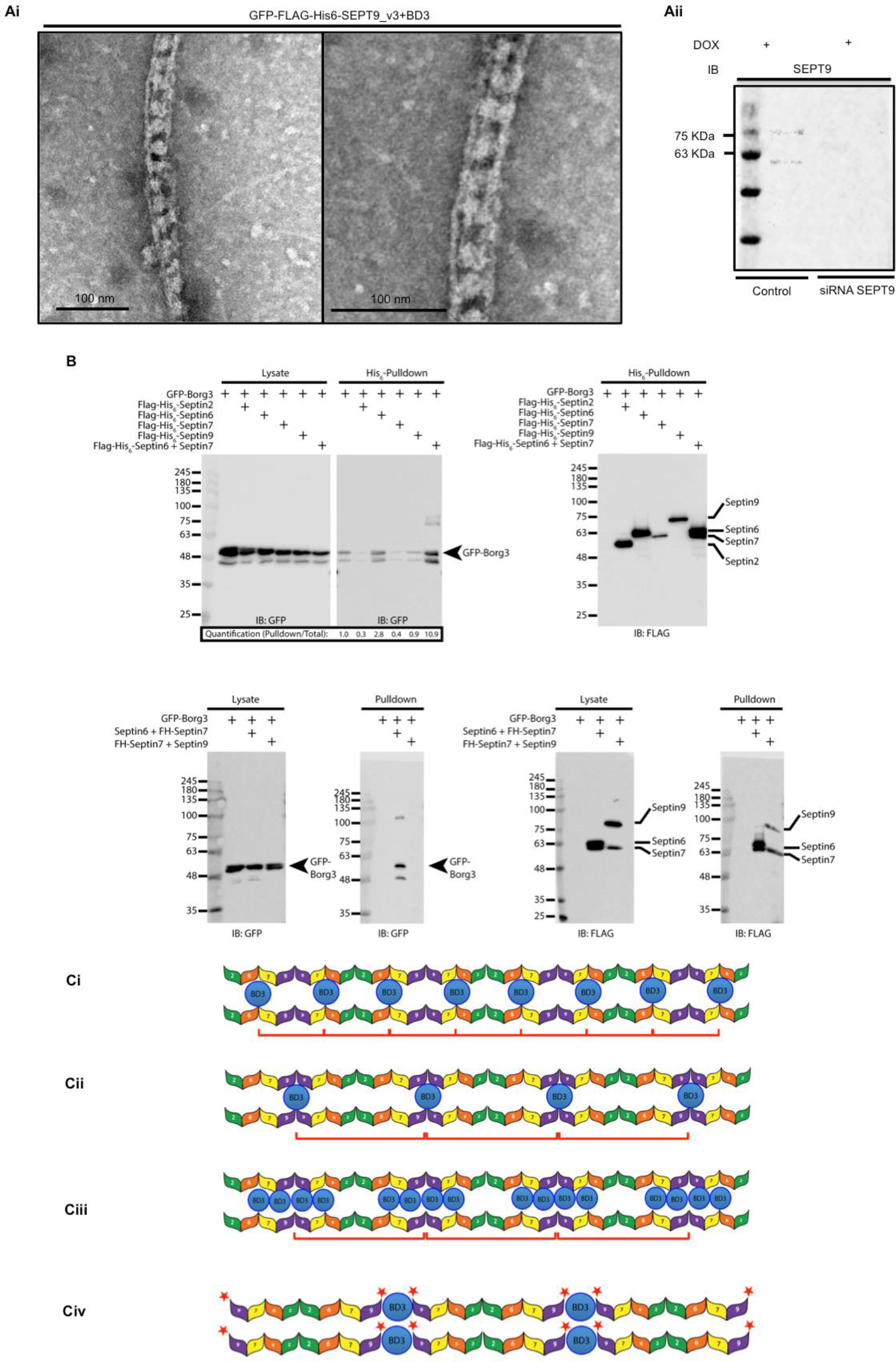
(Ai) Septin complexes isolated from GFP-FLAG-His6-SEPT9_v3 and BD3 domain was added to them and dialyzed against low salt buffer (50mM). Samples were examined with negative staining EM. (Aii) Immunoblotting with SEPT9 antibody that shows SEPT9 is depleted in the mChery-FLAG-SEPT7 cell line. (Bi) Individual septins (FLAG-His6-SEPT2, FLAG-His6-SEPT6, FLAG-His6-SEPT7, FLAG-His6-SEPT9) and septin dimer (FLAG-His6-SEPT6+ SEPT7) were expressed in the cells with GFP-Borg3. His6-pulldown was performed and immunoblotted with GFP antibody showing that the majority of GFP-Borg3 pulldown when FLAG-His6-SEPT6+ SEPT7 were expressed. (Bii) Septin dimmers (FLAG-His6-SEPT6+ SEPT7 and FLAG-His6-SEPT7+ SEPT9) expressed with GFP-Borg3 and His6-pulldown was performed. GFP-Borg3 pulldown with only FLAG-His6-SEPT6+ SEPT7 and not with FLAG-His6-SEPT7+ SEPT9. (Ci), (Cii) and (Ciii) Schematic representation of Borg binding sites on the mammalian septin octamers. (Ci) If Borgs were binding to SEPT7/SEPT6 the periodicity would be shorter than observed. Cii) If Borgs were to bind to SEPT9 periodicity would be similar to octamer length, but cross-bridges would be expected to be thin. (Ciii) Schematic representation of Borgs binding to both SEPT6/SEPT7 and SEPT9. Note: FFT measurements of more than 61cable-like structure in GFP-FLAG-His6-SEPT9_v5 reveal spacing of 35 nm, supporting models 2 or 3 (Civ) Schematic representation of BD3 domain binding to GFP-Flag-His6-SEPT9_ΔN-containing complexes. If BD3 domain links GFP-Flag-His6-SEPT9_ΔN, long filaments could be expected.

